# Clinical super-resolution computed tomography of bone microstructure: application in musculoskeletal and dental imaging

**DOI:** 10.1101/2023.06.28.544314

**Authors:** S.J.O. Rytky, A. Tiulpin, M.A.J. Finnilä, S.S. Karhula, A. Sipola, V. Kurttila, M. Valkealahti, P. Lehenkari, A. Joukainen, H. Kröger, R.K. Korhonen, S. Saarakkala, J. Niinimäki

**Author notes:** Corresponding author: Santeri J.O. Rytky. From the Research Unit of Health Sciences and Technology, University of Oulu, Oulu, Finland, POB 5000, FI-90014 Oulu, Finland. Data generated or analyzed during the study are available from the corresponding author by reasonable request.

## Abstract

**Objectives:** Clinical cone-beam computed tomography (CBCT) devices are limited to imaging features of half a millimeter in size. Hence, they do not allow clinical quantification of bone microstructure, which plays an important role in osteoarthritis, osteoporosis and fracture risk. For maxillofacial imaging, changes in small mineralized structures are important for dental, periodontal and ossicular chain diagnostics as well as treatment planning. Deep learning (DL)-based super-resolution (SR) models could allow for better evaluation of these microstructural details. In this study, we demonstrate a widely applicable method for increasing the spatial resolution of clinical CT images using DL, which only requires training on a limited set of data that are easy to acquire in a laboratory setting from e.g. cadaver knees. Our models are assessed rigorously for technical image quality, ability to predict bone microstructure, as well as clinical image quality of the knee, wrist, ankle and dentomaxillofacial region.

**Materials and methods:** Knee tissue blocks from five cadavers and six total knee replacement patients as well as 14 extracted teeth from eight patients were scanned using micro-computed tomography. The images were used as training data for the developed DL-based SR technique, inspired by previous studies on single-image SR. The technique was benchmarked with an *ex vivo* test set, consisting of 52 small osteochondral samples imaged with clinical and laboratory CT scanners, to quantify bone morphometric parameters. A commercially available quality assurance phantom was imaged with a clinical CT device, and the technical image quality was quantified with a modulation transfer function. To visually assess the clinical image quality, CBCT studies from wrist, knee, ankle, and maxillofacial region were enhanced with SR and contrasted to interpolated images. A dental radiologist and dental surgeon reviewed maxillofacial CBCT studies of nine patients and corresponding SR predictions.

**Results:** The SR models yielded a higher Pearson correlation to bone morphological parameters on the *ex vivo* test set compared to the use of a conventional image processing pipeline. The phantom analysis confirmed a higher spatial resolution on the images enhanced by the SR approach. A statistically significant increase of spatial resolution was seen in the third, fourth, and fifth line pair patterns. However, the predicted grayscale values of line pair patterns exceeded those of uniform areas. Musculoskeletal CBCT images showed more details on SR predictions compared to interpolation. Averaging predictions on orthogonal planes improved visual quality on perpendicular planes but could smear the details for morphometric analysis. SR in dental imaging allowed to visualize smaller mineralized structures in the maxillofacial region, however, some artifacts were observed near the crown of the teeth. The readers assessed mediocre overall scores in all categories for both CBCT and SR. Although not statistically significant, the dental radiologist slightly preferred the original CBCT images. The dental surgeon scored one of the SR models slightly higher compared to CBCT. The interrater variability κ was mostly low to fair. The source code (https://doi.org/10.5281/zenodo.8041943) and pretrained SR networks (https://doi.org/10.17632/4xvx4p9tzv.1) are publicly available.

**Conclusions:** Utilizing experimental laboratory imaging modalities in model training could allow pushing the spatial resolution limit beyond state-of-the-art clinical musculoskeletal and dental CBCT imaging. Implications of SR include higher patient throughput, more precise diagnostics, and disease interventions at an earlier state. However, the grayscale distribution of the images is modified, and the predictions are limited to depicting the mineralized structures rather than estimating density or tissue composition. Finally, while the musculoskeletal images showed promising results, a larger maxillofacial dataset would be recommended for training SR models in dental applications.

## INTRODUCTION

Image quality plays a pivotal role in assessing musculoskeletal and dental pathologies. The most common modalities in the field include magnetic resonance imaging (MRI), radiography, ultrasound, and computed tomography (CT)^1–3^. While MRI provides excellent soft tissue contrast and radiography is widely available, CT imaging is the superior method for imaging changes in bone^2, 4, 5^. Clinical cone-beam computed tomography (CBCT) imaging devices can achieve a voxel size of up to 100-200µm^3^ and are useful for detecting both orthopedic^6^ and dental pathologies^7^, joint trauma imaging^8^, and radiotherapy planning^9, 10^. For example, CBCT has been recognized as the recommended modality for assessing wrist fractures^8, 11^. despite the mentioned resolution, from the Nyquist’s theorem, the perceived *spatial resolution* is at least twice lower, and thus the visible clinical features in CBCT can only be of 500µm in size^12^. This, however, is not enough to observe bone microstructural changes.

Phantoms, that is, tissue-simulating test objects are scanned to calculate a modulation transfer function (MTF) and quantify the CT spatial resolution in a clinical setting^13, 14^. In practice, a series of line pair patterns^13^ or a high-contrast edge^15, 16^ can be used to estimate the MTF. The image quality of clinical devices is limited by multiple factors. The ones for X-ray imaging are radiation dose, motion, acquisition geometry, receptor size and the focal spot size of the beam. The resolution limit of clinical CT is roughly seven line pairs per centimeter^17^.

The bone microstructure is conventionally seen only with laboratory micro-computed tomography (µCT) devices. For measurement in a clinical setting, CBCT is the most promising modality^18^. As an example, bone microstructural changes are known to be associated with osteoarthritis severity^19^, and could be useful in the assessment of osteoporosis, bone strength and fracture risk^20, 21^. Detection of early osteoarthritis could facilitate earlier intervention, significantly reducing the socio-economic impact of the disease^22^. Karhula *et al*. have previously shown that bone subresolution features can be estimated with CBCT using texture analysis^23^. Individual quantitative parameters cannot be directly connected to local tissue changes but could be visible from high-quality images. Finally, dentomaxillofacial CBCT imaging requires high image quality for multiple indications. The trabecular bone microstructure is one of the key factors for dental implant planning^24^. Dental and periodontal diagnostics^12^, as well as assessment of ossicular chain and inner ear pathologies^25^, are all focused on assessing changes in tiny, mineralized structures.

One approach to increase image resolution, is to improve upon the reconstruction technique. Recent advancements include iterative-^26, 27^, model-based-^28^, and learned^29, 30^ reconstruction. However, these methods naturally require access to the raw CT projection images, access to which is typically restricted by the scanner’s manufacturer. Another method for upscaling, could simply rely on image interpolation combined with antialiasing. However, such techniques have difficulties in removing artefacts and blur from the approximated high-resolution images^31^.

Due to recent advancements in deep learning (DL), super-resolution (SR) methods can be used to predict impressive details from low-resolution images^32^. They are based on convolutional neural networks (CNN), that either modify the original input image or generate entirely new images from latent space. High- and low-resolution images are used in the training process with different approaches: unpaired training aims to match two datasets with different image quality without exact matches for each image^33, 34^. It is also possible to obtain only the high-resolution dataset and artificially distort the data to create matching low-resolution images^32^. Finally, the dataset could be collected using both low- and high-resolution imaging modalities and a subsequent co-registration. However, accurate co-registration is likely challenging in the case of highly distorted images.

Previously, SR has been used to increase MRI quality for the knee by Chaudhari et al^35, 36^. The authors thoroughly evaluate the performance of the SR method for visualizing cartilage morphometry and osteophytes. Brain MRI SR has also been assessed for clinical image quality^37^. The first SR studies for inner-ear CBCT have been introduced using generative adversarial networks^38^. Finally, µCT imaging and SR has been used to assess bone microstructure in a preclinical setting^39^. The DL methods are mainly criticized for their “black-box” nature and lack of interpretability. However, some deep learning SR algorithms are already on the market for CT^29, 40^ and MRI^37^. Thus, guidelines and recommendations for thorough clinical validation of such algorithms are needed. Before clinical use of SR, it would be crucial to ensure that the CNN predictions only increase the image quality and do not add new or remove existing pathological features from the images^41^.

In this study, we demonstrate how to develop widely applicable methods for increasing the spatial resolution of clinical CT images using DL, and how to properly validate the methods in several clinical domains. Our contributions are the following: (1) We assess the performance of SR methods for predicting 3D bone microstructure on independent data, quantifying the bone parameters. The technical image quality of the algorithm is assessed using phantom imaging and MTF analysis; (2) To show the versatility of the method, we enhance clinical CBCT images of knee, wrist, ankle and teeth, using models trained solely on a limited amount of preclinical data. The dental image quality is quantified in a reader study; (3) We release the pretrained SR networks and the source code, facilitating further development of the musculoskeletal and dental imaging field.

## MATERIALS AND METHODS

### Training data

The training data consists of twelve knee tissue block samples extracted from five healthy cadavers and six total knee arthroplasty (TKA) patients (Table 1). An overview of the i1mage data acquisition is in Figure 1. The sample harvesting was approved by the Ethical committee of Northern Ostrobothnia’s Hospital District (PPSHP 78/2013) and the Research Ethics Committee of the Northern Savo Hospital District (PSSHP 58/2013 & 134/2015). The tissue blocks are stored in phosphate-buffered saline after surgery, and subsequently imaged with a preclinical µCT scanner (Bruker Skyscan 1176; 80kV, 125µA, 26.7 µm voxel size). The images were reconstructed using the manufacturer’s software (NRecon, beam hardening and ring artefact corrections applied).

Furthermore, a total of fifteen human teeth were collected from nine patients with a tooth removal operation (Table 1, PPSHP 123/2021). The teeth were scanned using a laboratory desktop µCT scanner (Skyscan 1272, Bruker Inc., Kontich, Belgium; parameters: 100kV, 100µA 19.8µm voxel size, Cu 0.11mm filter). The reconstruction was conducted using the Nrecon software with beam hardening and ring artefact corrections applied. The reconstructions of fourteen extracted teeth from eight patients were used to provide further training data for the SR model in the case of dental CBCT. A tooth scan of one of the patients was excluded due to corrupted data in the µCT scan.

### Ex vivo test set

To provide the ground truth reference for bone microstructure prediction, we utilized a previously collected dataset^23^ consisting of 53 osteochondral samples from nine TKA patients and two deceased cadavers without an OA diagnosis (Table 1; ethical approval PPSHP 78/2013, PSSHP 58/2013 & 134/2015). The samples were imaged using two devices: a clinical extremity CBCT (Planmed Verity, Planmed Inc., Helsinki, Finland; parameters: 80kV, 12mA, 200µm voxel size, 20ms exposure time) and a laboratory desktop µCT scanner (Skyscan 1272, Bruker Inc., Kontich, Belgium; parameters: 50kV, 200µA 2.75µm voxel size, 2200ms exposure time, 0.5mm Al filter). The projection images were reconstructed with the corresponding manufacturer’s reconstruction software with a “standard” reconstruction filter applied for CBCT, and beam-hardening and ring artefact corrections were applied for µCT (Nrecon, v.1.6.10.4, Bruker microCT). The reconstructed volumes were coregistered to the same coordinate system using rigid transformations on the Bruker Dataviewer-software (version 1.5.4, Bruker microCT).

### Clinical images

The proposed method was further tested on clinical data acquired using the same Planmed Verity CBCT device (Table 1). The clinical dataset consists of one knee scan (50-year-old female; 96kV, 8mA, 200µm voxel size, 10s exposure time, “flat” reconstruction filter), one wrist scan (56-year-old female; 90kV, 6mA, 200µm voxel size, 6s exposure time, flat filter), and one ankle scan (34-year-old male; 96kV, 8mA, 400µm voxel size, 6s exposure time, flat filter). In the case of the knee and ankle, the imaging was done in the weight-bearing position. The participants are healthy volunteers, and the CBCT scans were acquired from the Oulu University Hospital digital research database. Finally, preoperative CBCT scans (Planmeca Promax; parameters: 120kV, 5-6mA, 200µm voxel size, 8s exposure time) were collected from the nine patients with tooth removal (ethical permission PPSHP 123/2021).

To validate the technical image quality, a commercially available CT quality assurance phantom (GE Healthcare, model no. 5128754) was imaged using a diagnostic CT device (GE Revolution Frontier; parameters: 120kV, 335mA, 730ms exposure time, 625µm pixel size, 5mm slice thickness, head filter).

**Table 1.** Dataset descriptions. Samples from both total knee arthroplasty patients and asymptomatic cadavers were used in the preclinical training and test sets. Different patients were included for training and testing. The test set characteristics are described in further detail by *Karhula et al* ^23^. Clinical studies were used to validate the method on realistic use cases.

| Preclinical datasets | # images | # samples (n) | # patients (N) |
| --- | --- | --- | --- |
| Knee tissue blocks | 220 544 | 12 | 11 |
| Extracted teeth | 45 540 | 14 | 8 |
| Ex vivo test set | 1 700 | 53 | 11 |
| Clinical studies |  |  |  |
| Wrist CBCT | 313 |  | 1 |
| Ankle CBCT | 219 |  | 1 |
| Knee CBCT | 471 | 1 | 1 |
| Dental CBCT | 3 352 |  | 9 |
| CT Quality assurance phantom | 6 |  | N/A |

**Figure 1.**
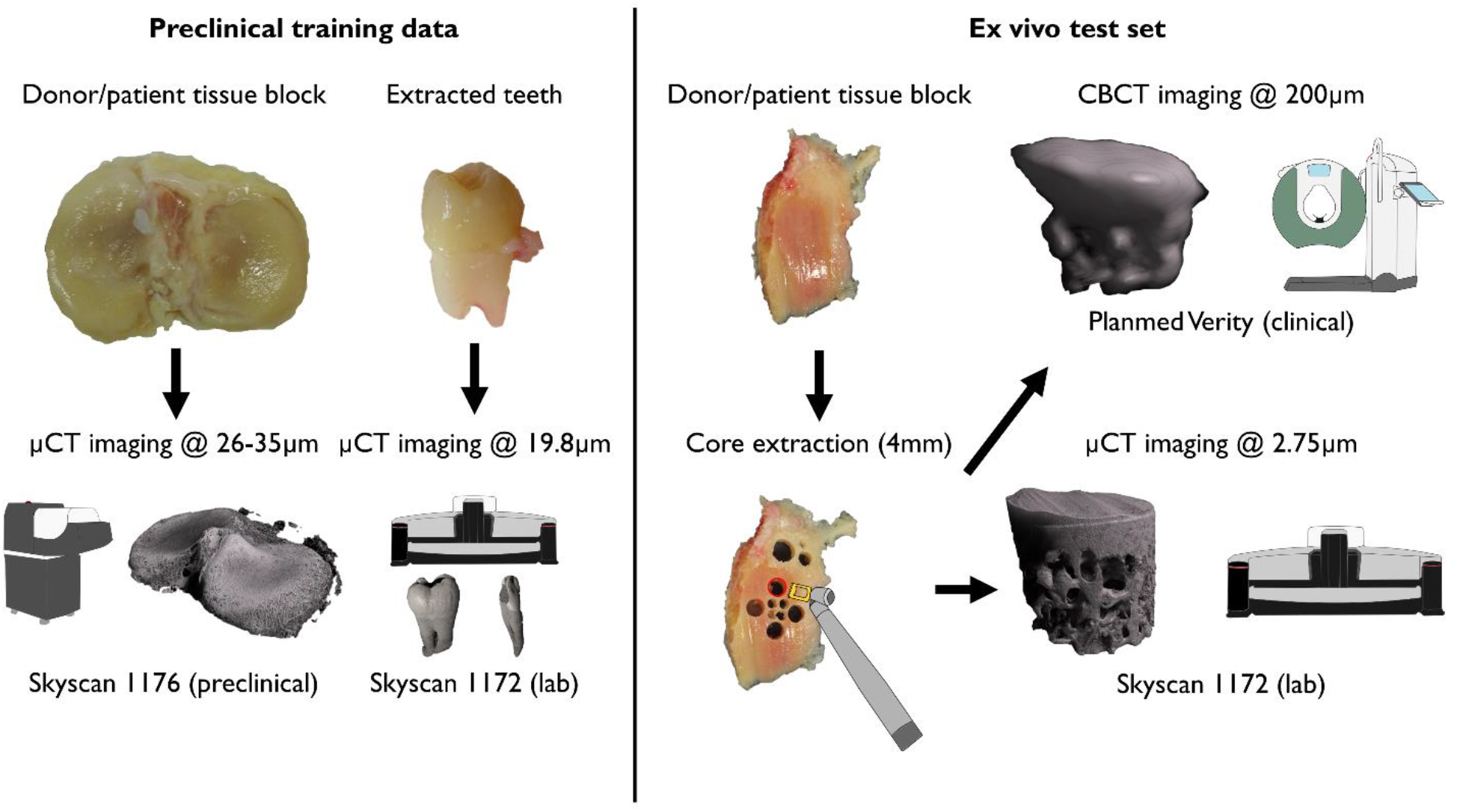
Training data and *ex vivo* test set acquisition. The full tissue blocks were scanned using a preclinical micro-computed tomography (µCT) scanner (Skyscan 1176). Extracted teeth were imaged using a desktop µCT (Skyscan 1272). Small 4mm osteochondral plugs were extracted and imaged both with the desktop µCT (Skyscan 1172) and a clinical extremity cone-beam CT (CBCT) system (Planmed Verity) to provide realistic low-and high-resolution references.

### Super-resolution model

The training data was created from the preclinical tissue blocks using interpolation. The three specific imaging resolutions used and the corresponding 4x magnifications were matched (200µm→50µm, 400µm→100µm, 488µm→122µm). To account for aliasing artefacts and simulate the lower imaging quality, Gaussian blurring (kernel size = 4, σ = 1) and median filtering (kernel size = 3) were applied after downscaling. The reconstructed image stacks were automatically divided into smaller 32x32x32 (input resolution) and 128x128x128 (target resolution) voxel patches suitable for training the SR models, resulting in thousands of training images (Table 1). The training data was augmented spatially using random rotations, translations and flips. Furthermore, brightness and contrast were randomly adjusted, and random blurring was added to augment the grayscale values. Finally, the input and target volumes were randomly cropped and padded to match the network input and output size (16x16→64x64 for 2D, 16x16x16→64x64x64 for 3D models). The augmentations were based on our previously published SOLT library (https://github.com/Oulu-IMEDS/solt) and modified to account for the varying input and target image size.

The model architecture was inspired by Johnson et al^42^, including four residual blocks (Figure 2, top). The transposed convolution layer was replaced by resize-convolution^43^. The model was designed to yield a magnification factor of four. To conduct the training process, we used an in-house developed Collagen framework (https://github.com/MIPT-Oulu/Collagen). We used three combinations of loss functions in the experiments: 1) The **baseline model** utilized mean squared error (MSE) and total variation (TV) as traditional pixel-wise losses, with respective weights of 0.8 and 0.2. 2) The **structure model** optimized the complement of the structure similarity index (SSIM), aiming to capture the bone microstructure. 3) The **visual model** combined mean absolute error (MAE), TV and perceptual loss (PL), aiming to provide the best perceptual quality, using weights of 0.1, 1.0 and 1.0, respectively. Features from a pretrained VGG16 model were used as the PL (Figure 2, bottom). The weights of the loss functions were chosen manually during the initial experiments of the study.

The models were trained using the Adam optimizer (parameters: α=0.0001, β=0.0001) for 50 epochs. The training was conducted under four-fold cross-validation, accounting for the patient ID during splits. During inference, the predictions were combined using a sliding window (16x16-pixel window with 8x8-pixel steps). A Gaussian kernel was applied to only focus the model predictions on the center of the tile, reducing the edge artefacts. To assess the performance of training, pixel-wise metrics (MSE, PSNR, SSIM) were calculated for the validation folds.

**Figure 2.**
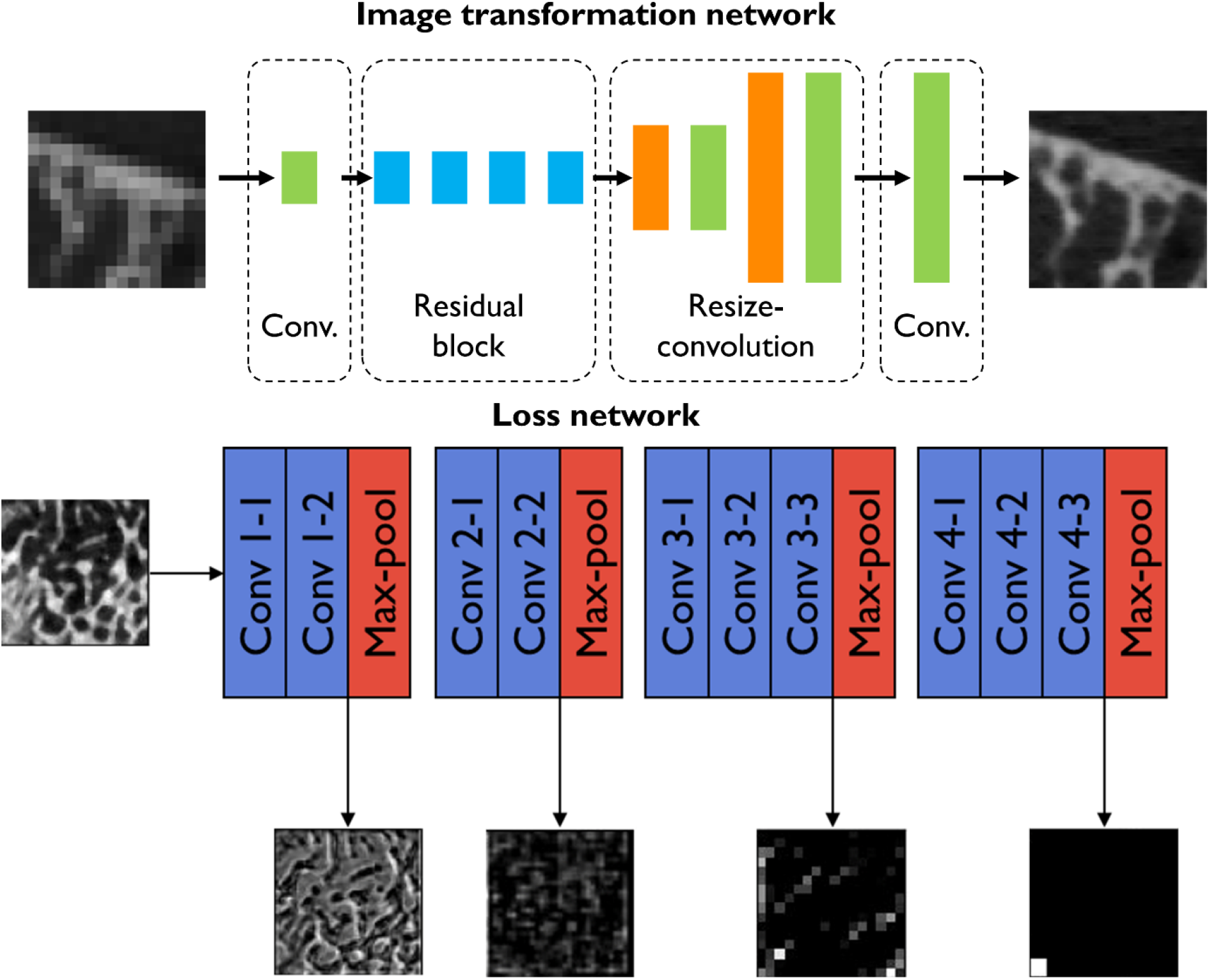
**Top:** The SR architecture used in the study. The architecture of Johnson et al. was modified by including resize-convolution layers instead of transposed convolutions. **Bottom:** The perceptual loss network was used in the visual model. Examples of perceptual loss network activations are shown for a trabecular bone reconstruction.

### Bone microstructure analysis

Morphological 3D parameters were quantified from the CBCT-imaged *ex vivo* test set, using conventional image processing, and SR. The true microstructure was analyzed using high-resolution µCT imaging. The volumes were binarized using the Otsu threshold ^44^. An ad-hoc Python script was used to calculate the recommended morphological parameters; bone volume fraction (BV/TV), trabecular thickness (Tb.Th), trabecular separation (Tb.Sp), and trabecular number (Tb.N)^45^. In the case of the 2D models, the parameters were assessed for the axial 2D predictions as well as an average of the predictions of the three orthogonal planes. To provide benchmark comparisons, tricubic interpolation, an image processing-based pipeline, and deep learning-based segmentations were used. The image processing pipeline included multiple subsequent filters prior to the binary thresholding (anisotropic diffusion, contrast stretching, median filter). The deep learning segmentation models consist of a ResNet-50 encoder with UNet and FPN decoders. Finally, the results were compared using Pearson correlation. The 95% confidence intervals were estimated for the models that are trained on multiple random seeds.

### Clinical validation images

To assess the technical image quality, the spatial resolution was quantified from the reconstructed phantom images and SR predictions. This was achieved by estimating the MTF using the six line pair patterns. The standard deviation was determined from a rectangular region-of-interest including each of the line pairs to provide a practical assessment of the function^13^. The frequency of 0.5 MTF (MTF_50%_) and 0.1 MTF (MTF_10%_), corresponding to a half-value and the limit of spatial resolution, are estimated from the graph.

To demonstrate the validity of the method in the clinical domain, we tested the models on multiple clinical imaging targets: ankle, knee, wrist and dental CBCT. The predictions and interpolated CBCT images were compared visually. The reconstructions were normalized and converted from 16-bit to 8-bit images. To save memory and computational time, small volumes of interest were selected from the wrist and the ankle (wrist = 6.3 x 6 x 3.7 cm, ankle = 6.6 x 6.3 x 4.8 cm). For the knee scan, the full joint was processed (10 x 10 x 10 cm, output size = 1884 x 1932 x 1988 voxels) on the Puhti supercomputer (https://research.csc.fi/csc-s-servers). For the ankle, a lower resolution is used, and another set of models is trained (400µm→100µm). In the case of knee, wrist and dental imaging, high-resolution models are used (200µm→50µm).

The predictions and interpolations from the preoperative dental CBCT scans were assessed in a blinded reader study by an experienced dental radiologist (Reader 1) and dental surgeon (Reader 2) to grade the level of diagnostic quality. The Likert scale was used to score the signal-to-noise ratio, anatomical conspicuity (periodontal ligament space), image quality, artifacts and diagnostic confidence of the images. The mean and standard deviation for the grades are reported and the inter-rater agreement is assessed using linearly-weighed Cohen’s Kappa (κ). Finally, two µCT scans of the extracted teeth are coregistered with the clinical scans to allow a further visual comparison (Dataviewer, v. 1.5.6.2).

## RESULTS

The conventional pixel-based performance metrics of training the 2D and 3D SR models on a 200µm→50µm resolution scale are summarized in Table 2. The 2D baseline model (trained with MSE+TV loss) yields the highest performance (MSE=0.0072±0.0002, PSNR=26.64±0.07, SSIM=0.812±0.003). The 2D structure and visual models as well as the 3D baseline model yield slightly higher errors. The lowest performance was with the 3D structure model (MSE=0.3±0.5, PSNR=15±15, SSIM=0.4±0.5).

**Table 2.** Results on the out-of-fold validation for the 200µm→50µm resolution models. Experiments with different combinations of loss functions are listed with a two-dimensional (2D) or volumetric (3D) model. The value for the standard error of mean is reported after the mean value.

| Models | Out-of-fold evaluation |  |  |
| --- | --- | --- | --- |
|  | MSE | PSNR | SSIM |
| Baseline 2D | **$0.0072 \pm 0.00003$** | **$26.64 \pm 0.014$** | **$0.812 \pm 0.0005$** |
| Structure 2D | $0.0084 \pm 0.0001$ | $25.5 \pm 0.05$ | $0.776 \pm 0.006$ |
| Visual 2D | $0.015 \pm 0.007$ | $25 \pm 1.3$ | $0.7 \pm 0.06$ |
| Baseline 3D | $0.0068 \pm 0.0001$ | $24.8 \pm 0.05$ | $0.691 \pm 0.002$ |
| Structure 3D | $0.3 \pm 0.11$ | $15 \pm 3.5$ | $0.4 \pm 0.1$ |
| Visual 3D | $0.02 \pm 0.005$ | $19 \pm 1$ | $0.4 \pm 0.04$ |

MSE=mean squared error, PSNR=peak signal-to-noise ratio, SSIM=structure similarity index

### Ex vivo test set: prediction of bone microstructure

The trained models were applied to the *ex vivo* test set to assess the performance of predicting the bone microstructure on unseen data (Table 3, Figure 3). The 2D structure model yields the highest results (r_BVTV_ = 0.817±0.005) and outperforms the interpolation (r_BVTV_ = 0.64) and conventional segmentation pipeline (r_BVTV_ = 0.67). The deep learning segmentation models did not converge on the dataset and all images were classified as background.

**Table 3.** Quantification of the bone parameters. Predictions from each model were binarized and the bone parameters were compared to the micro-computed tomography (µCT) ground truth. The values indicate Pearson correlations and the respective 95% confidence intervals. The highest correlation on each parameter is bolded. The deep learning models (ResNet-34 with UNet and ResNet-34 with FPN) did not generalize to the ex vivo test set.

| Models |  | Bone parameters |  |  |  |
| --- | --- | --- | --- | --- | --- |
|  | Averaging | BV/TV | Tb.Th | Tb.Sp | Tb.N |
| Interpolation |  | 0.64 | 0.34 | 0.59 | -0.4 |
| Conventional segmentation |  | 0.67 | 0.42 | 0.50 | -0.63 |
| Baseline 2D | No | 0.736±0.006 | 0.404±0.008 | 0.694±0.004 | -0.514±0.007 |
| | Yes | 0.665±0.003 | 0.336±0.003 | 0.608±0.006 | -0.458±0.0001 |
| Structure 2D | No | **0.817±0.005** | **0.53±0.02** | **0.756±0.009** | **-0.489±0.007** |
| | Yes | 0.731±0.007 | 0.436±0.006 | 0.613±0.010 | -0.41±0.02 |
| Visual 2D | No | 0.758±0.012 | 0.453±0.011 | 0.70±0.02 | -0.57±0.02 |
| | Yes | 0.674±0.004 | 0.340±0.009 | 0.609±0.011 | -0.5±0.02 |
| Baseline 3D | | 0.654±0.010 | 0.33±0.03 | 0.63±0.011 | -0.34±0.03 |
| Structure 3D | | 0±1.3 | 0±0.6 | 0±1.1 | -0.5±0.4 |
| Visual 3D | | 0.69±0.04 | 0.39±0.05 | 0.6±0.07 | -0.34±0.09 |

BV/TV=bone volume fraction, Tb.Th=trabecular thickness, Tb.Sp=trabecular separation, Tb.N=trabecular number

**Figure 3.**
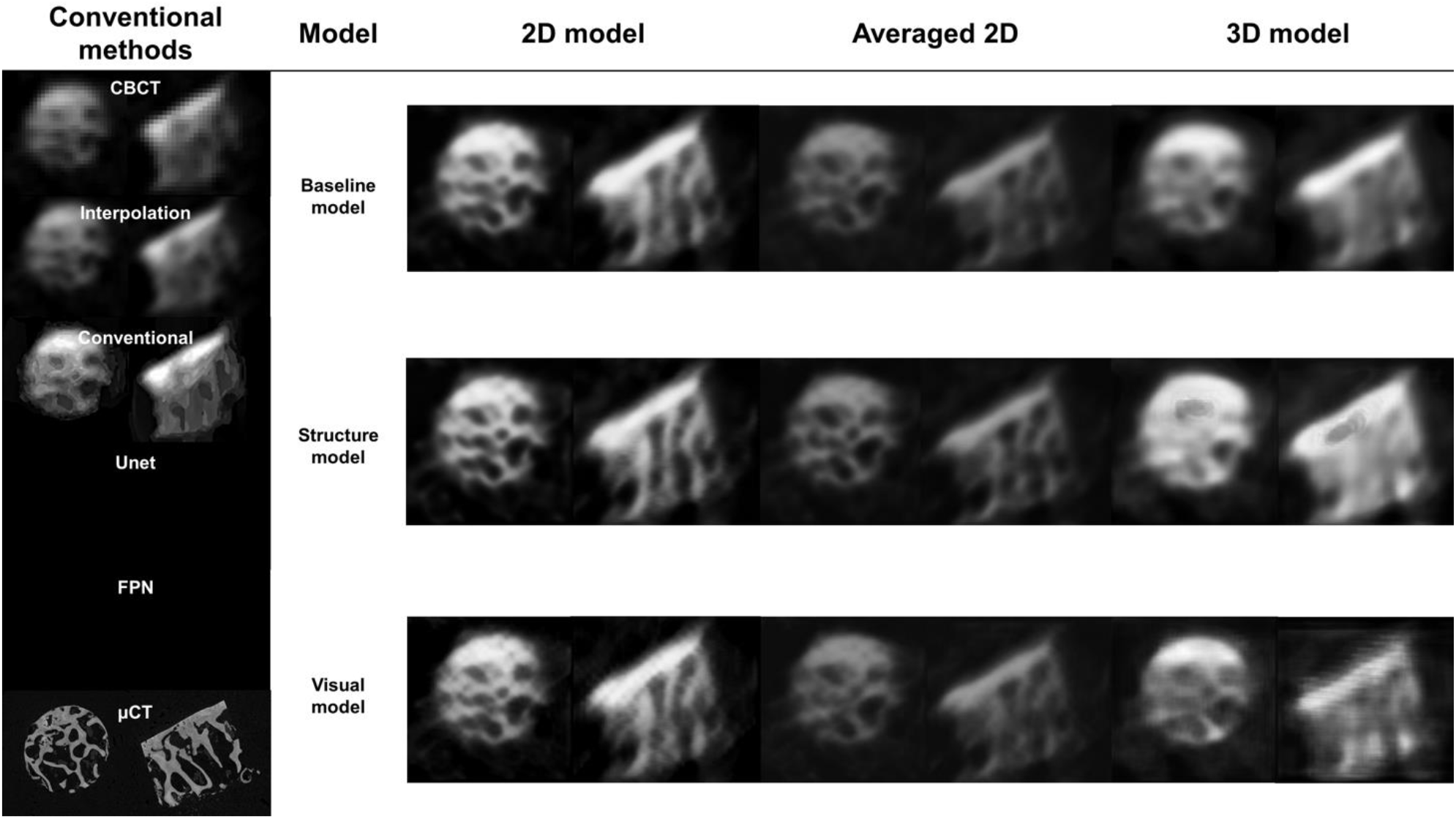
Comparison of conventional image quality improvement and super-resolution (SR) predictions on the osteochondral samples. The clearest structural definition is seen on the 2D models without averaging the three orthogonal planes. Deep learning segmentation was also tested, but all the samples were fully predicted as background (models did not generalize to the highly different test set).

### Technical image quality

The technical image quality was determined by comparing interpolated and predicted clinical CT images from a quality assurance phantom. The fifth line pair pattern at 8.3 line pair per cm frequency can be visually resolved from the SR predictions but not from the interpolated image (Figure 4a). Furthermore, the MTFs suggest a higher image quality in the predictions at the 4-8 line pairs per cm frequency range. An increase of 0.2 is seen between 5-6 line pairs per cm (Figure 4b). Based on the estimated MTF curves, the interpolated CT images reach MTF_50%_ and MTF_10%_ at roughly 3.5 and 7.0 line pairs per cm, respectively. The MTF curves from the SR models reach the MTF_50%_ and MTF_10%_ values later, at 5.0 and 8.0 line pairs per cm. Standardization based on plexiglass and water grayscale values was not feasible for the SR models (Supplementary Figure 1).

**Figure 4.**
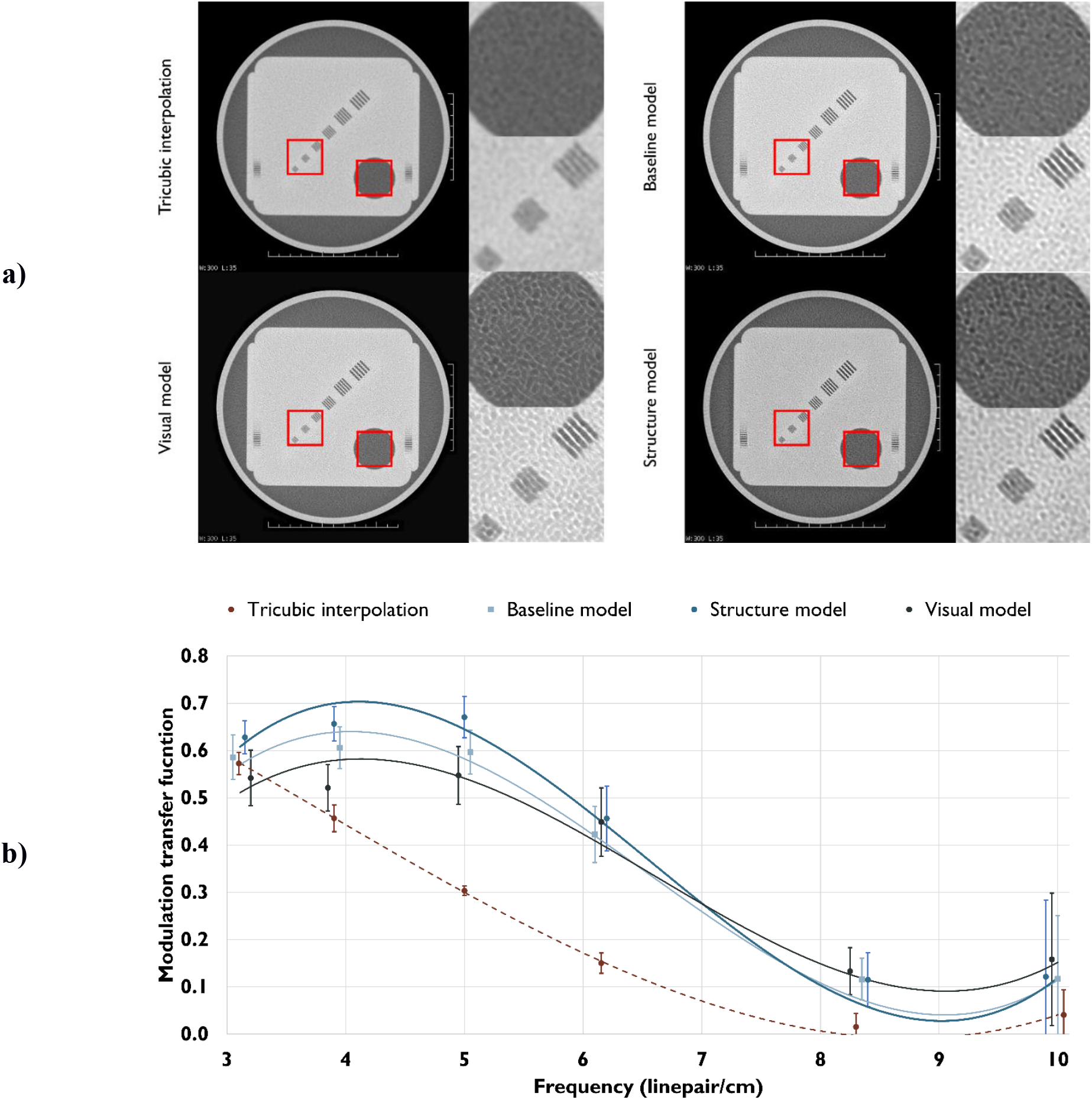
A clinical CT scan of a commercially available quality assurance phantom, with the corresponding interpolations and super-resolution (SR) predictions (a). Using the SR models, another set of line pairs can be distinguished from the CT slices. However, the perpendicular plane resolution is less improved. This can be seen as the number of diagonal lines on the edge of the phantom (that are averaged from multiple different depths) is not decreasing. The modulation transfer functions (MTF, b) show that all the SR models provide an increase in spatial resolution. The 95% confidence intervals are shown for each MTF measurement. Rough trendlines of the MTFs are shown with a third-order polynomial fit.

### Clinical image quality on musculoskeletal application

An overview of the proposed SR method and an example of wrist SR are presented in Figure 5. A volume-of-interest in the wrist joint was passed through the model to reduce the computational time. The computation on all three planes took roughly one hour on two graphical processing units (Nvidia GeForce GTX 1080 Ti). More structural details are visible in the prediction, but the cortical bone is visually too porous when compared to the original CBCT image. We also tested whether the inclusion of teeth images in training data changed the appearance, but only small differences were observed (Supplementary Figure 2) compared to the original training setup. In the case of knee CBCT, a large volume was processed on the Puhti supercomputer. The 2D models were compared to the interpolation and conventional image processing pipeline (Figure 6). The structural details were visually highlighted the best on the results from the baseline and structure models. The visual model created a flickering artefact in noisy and unclear regions of the tissue (Supplementary Video 1).

**Figure 5.**
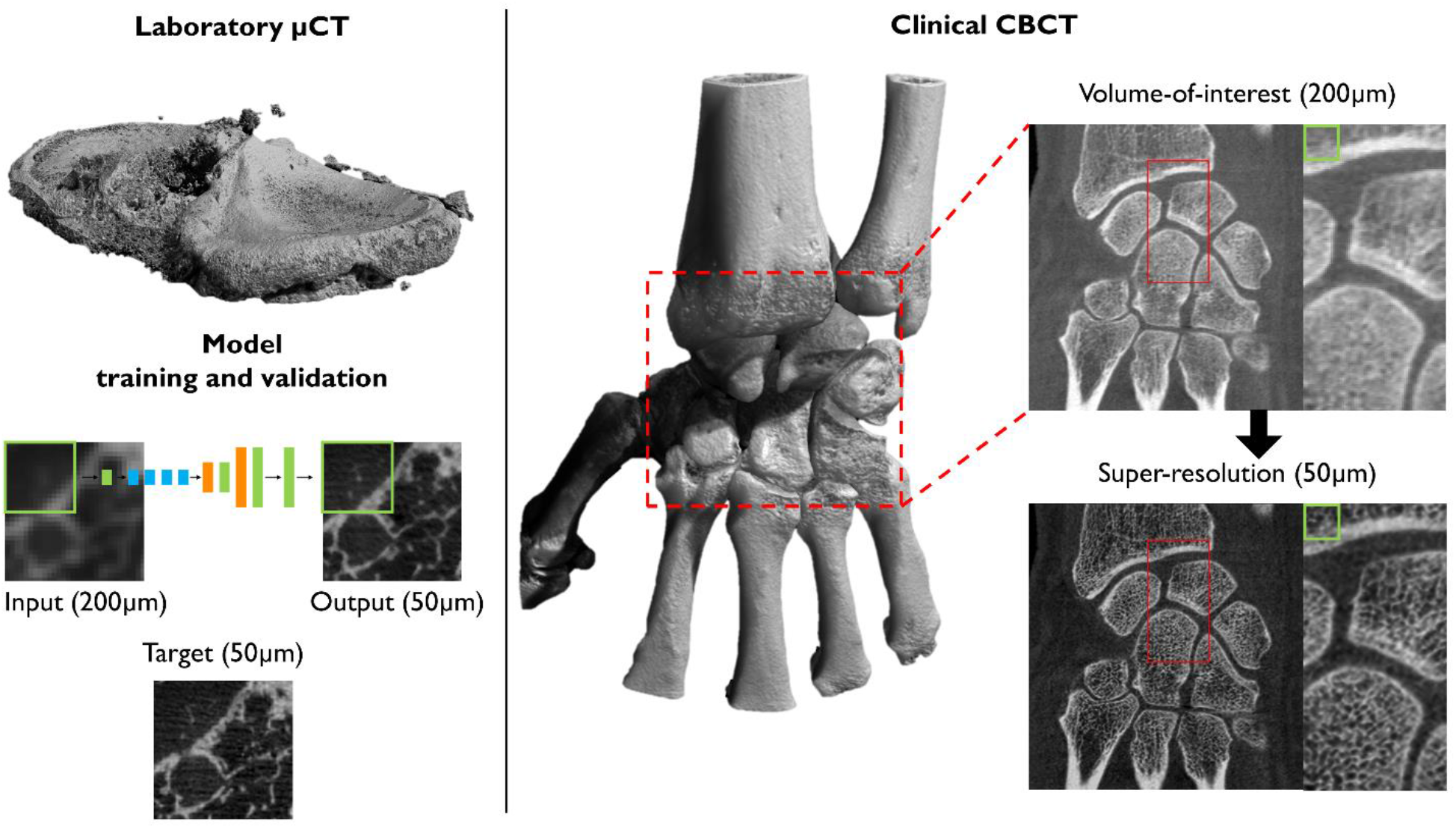
Overview of the proposed super-resolution (SR) method. Tissue blocks are scanned with micro-computed tomography (µCT) and used to train the model (left). The trained model can be utilized for clinical cone-beam CT (CBCT) images using a patch-by-patch sliding window, the size of one patch is depicted with a green rectangle. In this case, predictions from all orthogonal planes were averaged.

**Figure 6.**
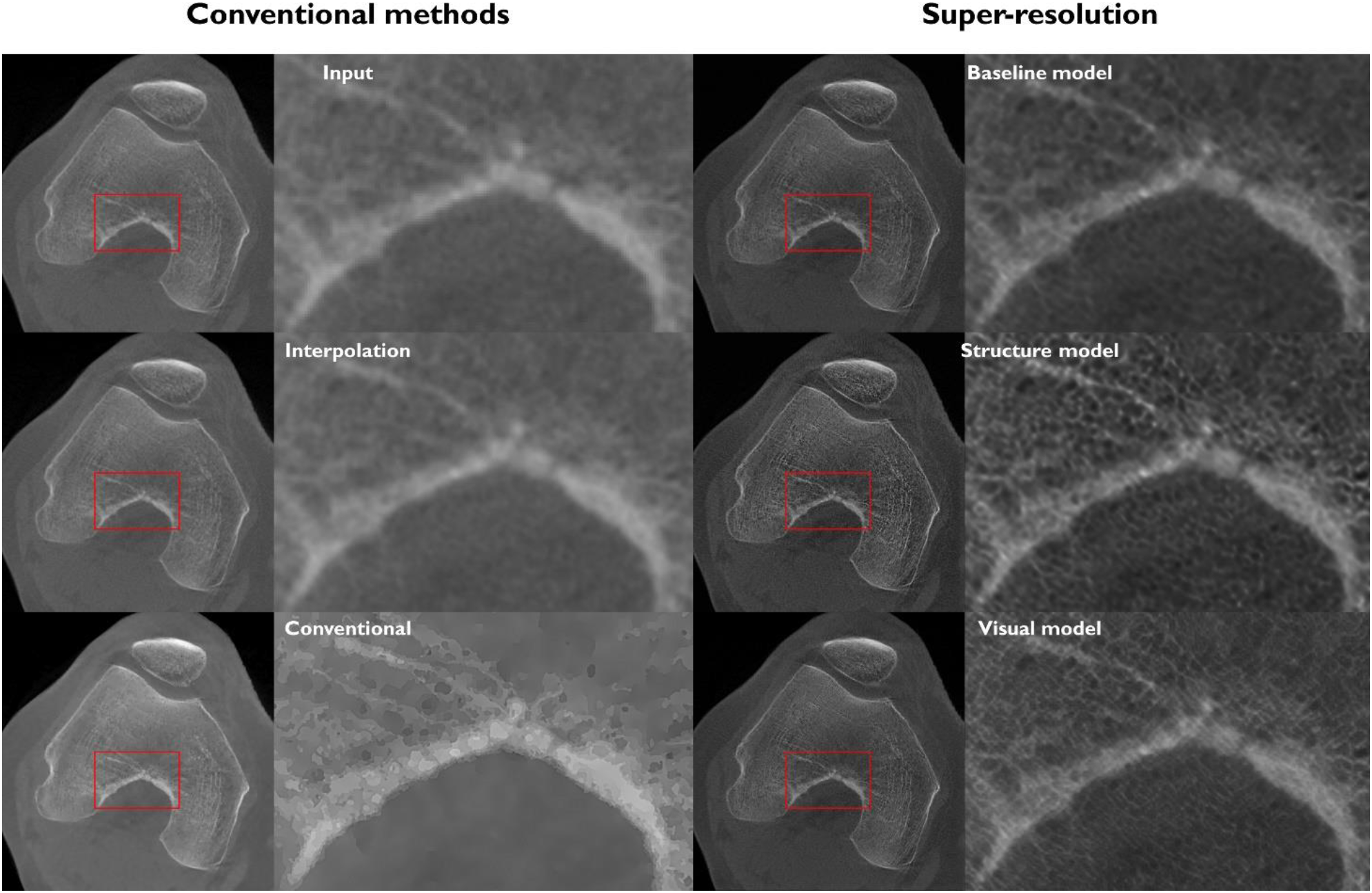
Comparison of conventional image quality improvement and super-resolution (SR) predictions on clinical scans of the knee joint. Predictions were conducted for the full joint; magnifications are shown to allow for a better visual comparison.

The ankle CBCT images were visually compared to interpolation, conventional image processing pipeline, as well as 2D and 3D predictions (Figure 7). The 2D models show reduced noise and slightly more details compared to the conventional methods. The most clearly visible structures were yielded by the structure model. None of the 3D models converged to a solution with sufficient image quality. This led to noisy prediction images, highlighting only the edges of the bones.

**Figure 7.**
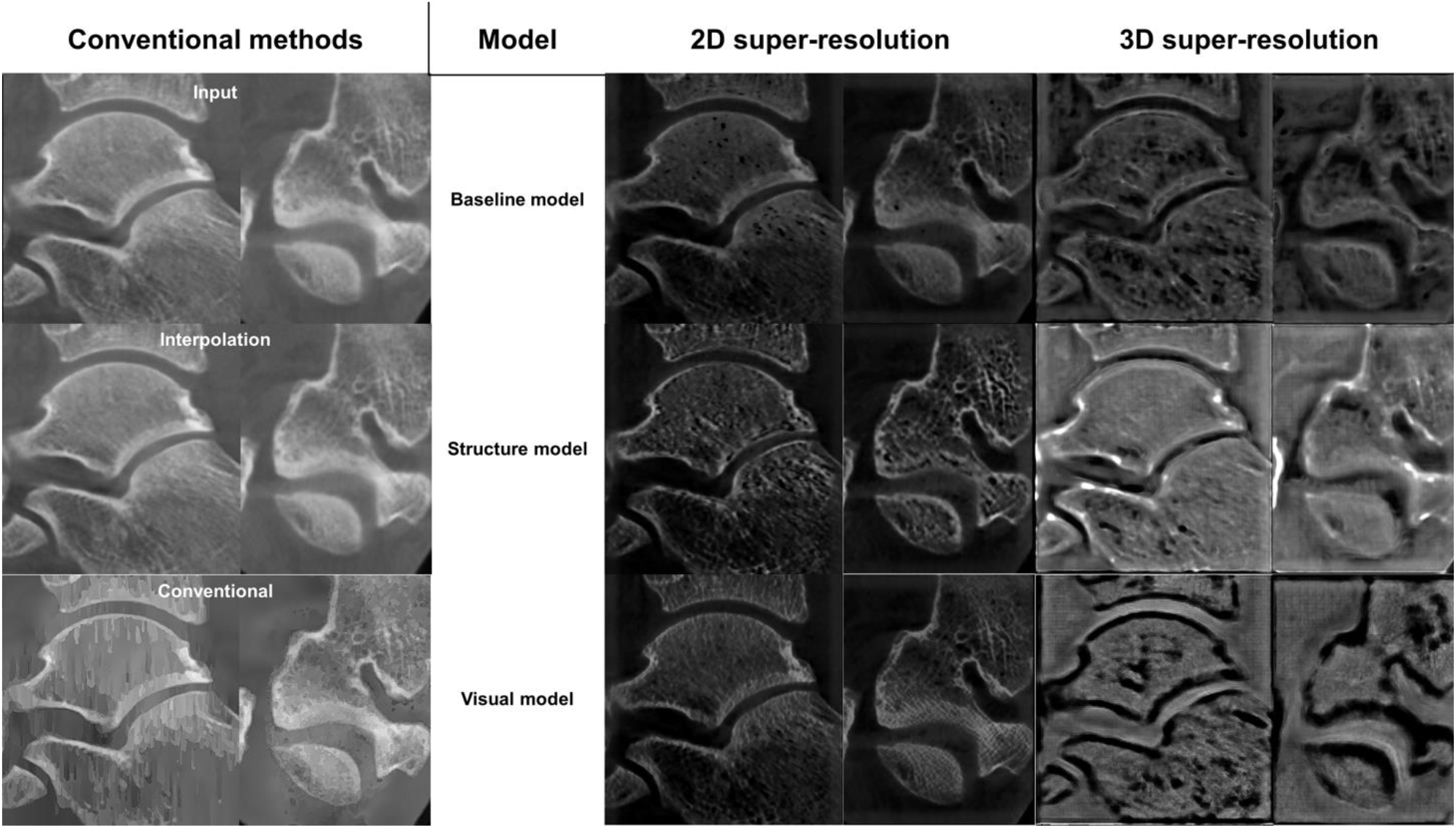
Comparison of conventional image quality improvement and super-resolution (SR) predictions on clinical scans of the ankle joint. The 3D adaptation of the SR models did not converge and provided noisy results.

### Clinical image quality on dental application

An example of SR prediction on maxillofacial CBCT is shown in Figure 8. In this case, the teeth of the patient were not used in training the SR model. A comparison of CBCT, SR and µCT of extracted teeth from two other patients is illustrated in Supplementary Video 2. Small structures are better highlighted on the SR images compared to the original CBCT, and a previously unseen gap can be seen in the lamina dura next to the tooth that was removed from patient one (indicated with a red arrow). We noted artefacts from the SR algorithms especially within the enamel. The results of the reader study are described in Table 4. When accounting for Bonferroni correction, no significant differences were observed for scores of Reader 1, although a slight trend of higher scores towards the interpolated images was observed. Reader 2 scored higher signal-to-noise ratio, anatomical conspicuity, image quality and diagnostic confidence for the baseline model compared to interpolation. The inter-rater agreement was slight (0.0-0.2) or fair (0.2-0.4), yet a substantial agreement was found for signal-to-noise ratio (0.64, visual model) and artifacts (0.80, baseline model).

**Figure 8.**
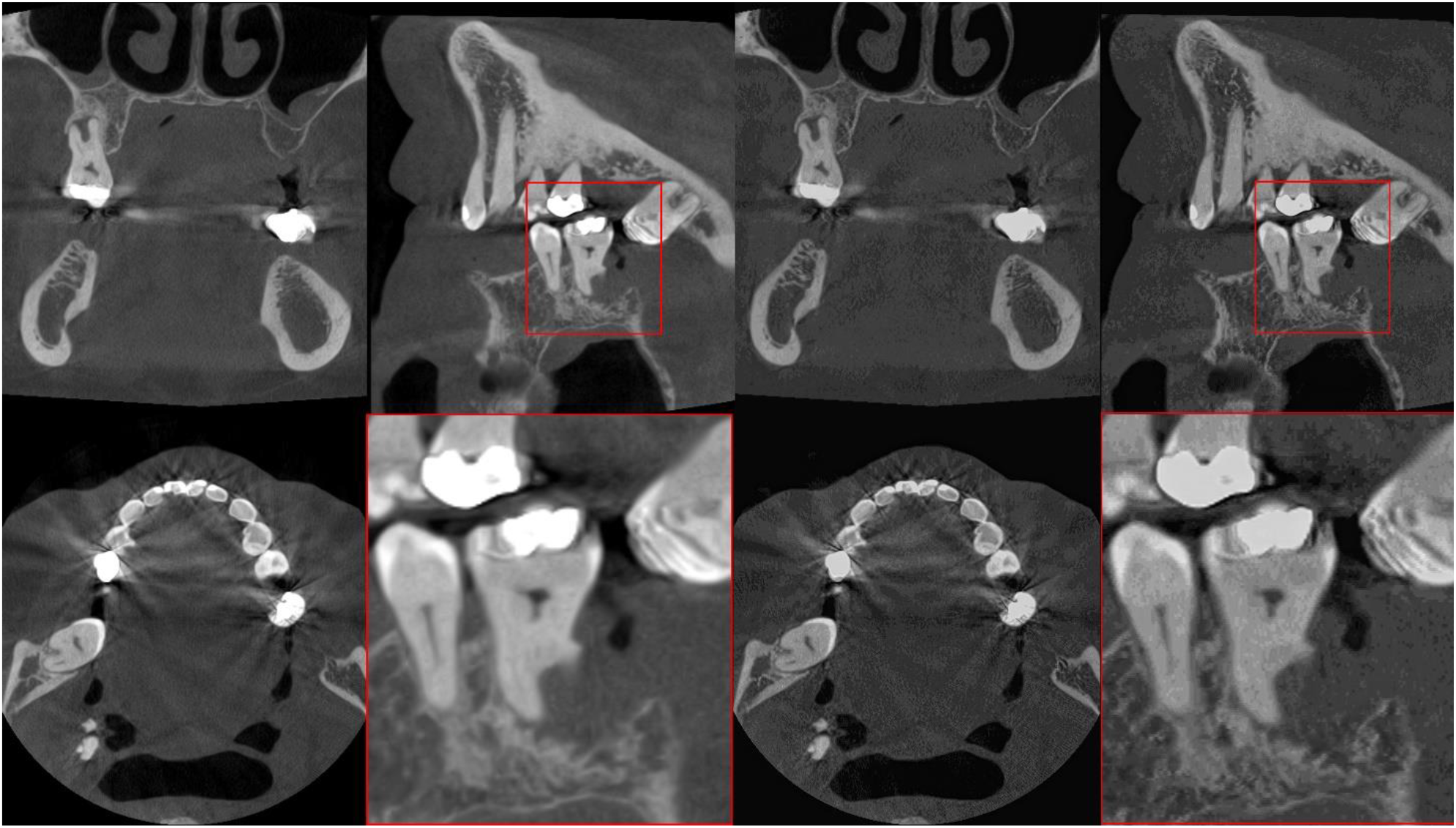
Examples maxillofacial cone-beam CT images (left) and corresponding super-resolution predictions (right). Predictions are shown from the structure model, without averaging the orthogonal planes.

**Table 4.** Blinded reader qualitative assessments. The mean and standard deviation of the scores are given for each category. The inter-reader agreement was assessed using Cohen’s Kappa (κ) with 95% confidence intervals. Statistical significance for differences between interpolation and super-resolution (SR) was assessed using the Wilcoxon Signed Rank test (Bonferroni corrected for three comparisons) and is indicated with an asterisk (*).

| Model | Score (Mean+SD) |  |  |  |  | Overall average |
| --- | --- | --- | --- | --- | --- | --- |
|  | Signal-to-noise ratio | Anatomical conspicuity | Image quality | Artifacts | Diagnostic confidence |  |
| Reader 1 |  |  |  |  |  |  |
| Interpolation | 2.4±0.7 | 2.9±0.6 | 2.8±0.4 | 2.7±0.5 | 2.8±0.4 | 2.7±0.5 |
| Baseline model | 2.2±0.4 | 2.7±0.5 | 2.6±0.5 | 2.7±0.5 | 2.7±0.5 | 2.6±0.5 |
| Structure model | 1.9±0.8 | 2.4±0.5 | 2.4±0.5 | 2.2±0.4 | 2.4±0.5 | 2.3±0.6 |
| Visual model | 2.1±0.3 | 2.4±0.5 | 2.3±0.5 | 2.3±0.5 | 2.7±0.5 | 2.4±0.5 |
| Reader 2 |  |  |  |  |  |  |
| Interpolation | 1.4±0.7 | 2.1±0.8 | 1.8±0.7 | 2.2±1.1 | 1.8±1.0 | 1.9±0.9 |
| Baseline model | 2.4±0.5* | 2.8±0.7* | 2.8±0.7* | 2.8±0.7 | 2.9±0.6* | 2.7±0.6 |
| Structure model | 1.4±0.5 | 2.0±0.5 | 1.8±0.7 | 2.1±1.1 | 1.9±0.8 | 1.8±0.7 |
| Visual model | 2.0±0.5 | 2.2±0.4 | 2.0±0.5 | 2.1±0.6 | 2.1±0.6 | 2.1±0.5 |
| Agreement (κ) |  |  |  |  |  |  |
| Interpolation | 0.147 | 0.241 | 0.047 | 0.077 | 0.039 |  |
| Baseline model | 0.526 | 0.400 | 0.250 | 0.800 | 0.143 |  |
| Structure model | 0.400 | 0.217 | 0.156 | 0.087 | 0.031 |  |
| Visual model | 0.640 | 0.053 | 0.308 | 0.143 | 0.211 |  |
| 95% CI |  |  |  |  |  |  |
| Interpolation | (0.108-0.186) | (0.224-0.258) | (0.018-0.076) | (0.048-0.106) | (0.006-0.071) |  |
| Baseline model | (0.518-0.534) | (0.389-0.411) | (0.238-0.262) | (0.793-0.807) | (0.132-0.153) |  |
| Structure model | (0.379-0.421) | (0.202-0.233) | (0.135-0.178) | (0.060-0.114) | (0.010-0.051) |  |
| Visual model | (0.636-0.644) | (0.043-0.062) | (0.298-0.317) | (0.132-0.153) | (0.196-0.225) |  |

CI=confidence interval, \* $p < 0.05$

## DISCUSSION

In this study, we presented a deep learning-based super-resolution method to increase medical CBCT image quality in musculoskeletal and dental imaging domains. The predictions were assessed using conventional image metrics, bone microstructure assessment, as well as through multiple experiments for clinical data. The technical increase in spatial resolution was quantified using a quality assurance phantom. Finally, the method was tested on clinical CBCT images of the wrist, knee, ankle and maxillofacial region. The validation experiments are completely independent of the training process. This simulates deploying a method developed on laboratory data in the clinical environment, which we consider one of the key strengths in this study. The source code of the project is published on GitHub (https://doi.org/10.5281/zenodo.8041943) and the pretrained models used for dental SR predictions are available on Mendeley Data (https://doi.org/10.17632/4xvx4p9tzv.1).

The out-of-fold validation results (Table 2) suggest that the 2D baseline model performs best and that the 3D models yield the lowest performance. The analysis is based on traditional pixel-wise comparisons to high-resolution images. However, the analysis of osteochondral *ex vivo* samples shows that the 2D structure model is the best for predicting microstructural bone details (r_BVTV_ = 0.817±0.005). Furthermore, averaging the prediction on three orthogonal planes did not improve the result. Likely, averaging the 2D predictions that do not account for adjacent slices causes smearing of the details, resulting in a lower correlation at least in the studied small four-millimeter samples. Finally, we would like to note that we also trained UNet and FPN segmentation models to predict the bone microstructure, but the models did not generalize from the training on the tissue blocks to the *ex vivo* test set. Thus, we hypothesize that the SR method is more resistant to domain shift. This is further supported by the multiple of applications presented using the same training data.

The results of the quality assurance phantom analysis suggested that the SR models increase CT spatial resolution, both visually and quantitatively. Importantly, we also noticed that the models heavily modified the grayscale distribution of the scan, and the values on the line pair pattern were exceeding those in the uniform areas of the phantom. This eventually led us to scale the MTF curves, based on the maximum intensity of the scan (Supplementary Figure 1). Importantly, the quantitative Hounsfield unit values are lost after processing, and the resulting prediction only describes the bone structure, not density or material composition. This is a potential limitation of patch-based super-resolution but could be alleviated in the future using wider dynamic range of training images or more complex SR models.

The experiments on the wrist, knee, ankle and maxillofacial region reveal that the models generalize very well on different anatomical regions, although some regions of cortical bone there is sudden increase in porosity, especially in the wrist images. This is likely a result of having a high amount of trabecular bone in the training data. However, this was not confirmed in Supplementary Figure 2, as there were no major differences in the images. In the maxillofacial region, our initial experiments included multiple artefacts near teeth, when using only the knee tissue blocks in training. Averaging the predictions in three orthogonal planes preserves the structure better in the perpendicular plane but might smear the details in case of morphological analysis. This is also supported by the Supplementary Video 1, where a flickering artefact is seen on the sagittal plane.

The reader study resulted in quite modest scores for both interpolated images and SR predictions. A slight preference for interpolated images was observed for the scores of Reader 1, and Reader 2 scored the Baseline model slightly higher compared to other models or interpolation. The low overall scores are likely due to the fact that the high dynamic range of the original 12-bit CBCT images is lost. This could be potentially alleviated in the future by training the models on a higher dynamic range rather than the conventional eight bits which would also better allow studying HU values of model output. Also, the volume of extracted teeth is very small, and the current dataset is not optimal for training SR models for dental images.

While promising maxillofacial images show that the small, mineralized structures are better seen on the SR predictions, and even previously unseen pathologies might be revealed (Supplementary Video 2). However, we also noted definite artefacts within the enamel which could be confused for caries lesions. A more specialized training dataset would be crucial to alleviate such issues. Indeed, we hypothesize that the best results would be obtained using a dataset with preclinical scans of entire cadaveric jawlines and soft tissues. Even more readily available animal models, such as pig maxillofacial tissue, could be considered to provide the SR models examples closer to the target distribution.

In medical diagnotics, it is imperative that the SR models do not induce biases from the training set and remove or add new diagnostic features to the predicted high-resolution images^41^. Upscaling the images poses a serious theoretical problem: multiple visually distinct high-resolution images can downscale to the same low-resolution image^46^. This serious limitation warrants thorough validation experiments before SR can be utilized in the clinical environment. This would be an excellent area for future studies, where predictions of healthy tissue and small fractures or other pathological conditions could be analyzed in more detail. We would hypothesize that models that generate entirely new images from a latent space, such as generative adversarial networks, could have a higher risk of “hallucinating” nonexistent pathological features, whereas a traditional CNN is more limited to modifying the original image, even though it is upscaled from low-resolution.

This study has several limitations. First, the best-performing 2D models did not account for changes in the perpendicular plane. An interesting future methodological improvement could include using a three-channel input image, including the adjacent slices. Second, most of the clinical comparisons presented in this study are restricted to qualitative or semiquantitative analysis. There are many studies where multiple radiologist readers assess the diagnostic image quality blindly from the SR and comparison images to show the increase in performance^36, 37, 47, 48^. We would argue that the ratings provided by the radiologists are also somewhat subjective, and the true ground-truth information cannot be obtained in clinical studies without a subsequent tissue sample extraction. Third, the weights of the individual loss functions were chosen manually during the early experiments of this study. These should be ideally chosen using an ablation study or hyperparameter optimization. Finally, the SR prediction was conducted as post-processing rather than by directly reconstructing the projection images using deep learning. Indeed, the first CT vendors have already released reconstruction methods based on deep learning^29, 40^. As the projection data is often unavailable to the end user, nonlinear CNN-based methods that work in the reconstruction domain could be more easily added, as an additional component to any CT scanner.

The deep-learning-enhanced medical images could have a high impact on the medical domain. The implications for the technology include higher patient throughput, more precise diagnostics, and disease interventions at an earlier state. The proposed SR can be directly applied to the existing clinical scans in the reconstruction domain and could thus have quality enhancement potential for routine hospital pipelines. Integration of SR methods in the hospital environment could facilitate a higher throughput, reducing the time radiologist needs to reach a diagnosis on difficult cases as well as mitigating uncertainty in the diagnostic process. Radiologists could use the SR as an advanced “zoom” feature, analogous to pathologist changing the objective on a microscope. Training the models on laboratory data allows for pushing the spatial resolution limit further than what the clinical radiation doses or even the current CT technology would otherwise allow. Alternatively, the current image quality could be maintained with a lower dose which could increase the accessibility of CBCT and allow earlier diagnostic intervention.

## ACKNOWLEDGEMENTS

The financial support of the Academy of Finland (grants no. 268378, and 303786); Sigrid Juselius Foundation; European Research Council under the European Union’s Seventh Framework Programme (FP/2007-2013)/ERC Grant Agreement no. 336267; 6GESS Profiling Research Programme of the University of Oulu (Academy of Finland project 336449); Instumentarium Science Foundation (grant no. 200058); Finnish Cultural Foundation (grant no. 00200953); KAUTE foundation, and the strategic funding of University of Oulu are acknowledged. The authors wish to acknowledge CSC – IT Center for Science, Finland, for computational resources.

## AUTHOR CONTRIBUTIONS

Conception and design: SJOR, AT, MAJF, SSK, SS, JN.

Development of the pipeline: SJOR, AT.

Data analysis: SJOR.

Data acquisition: SJOR, MAJF, SSK, AS, VK, MV, PL, AJ, HK, RK.

Drafting the manuscript: SJOR.

Critical revision for important intellectual content and approval of the manuscript: all authors.

## ROLE OF THE FUNDING SOURCES

Funding sources are not associated with the scientific contents of the study.

## COMPETING INTERESTS

The authors report no conflicts of interest.

## SUPPLEMENTARY DATA

**Supplementary Figure 1.**
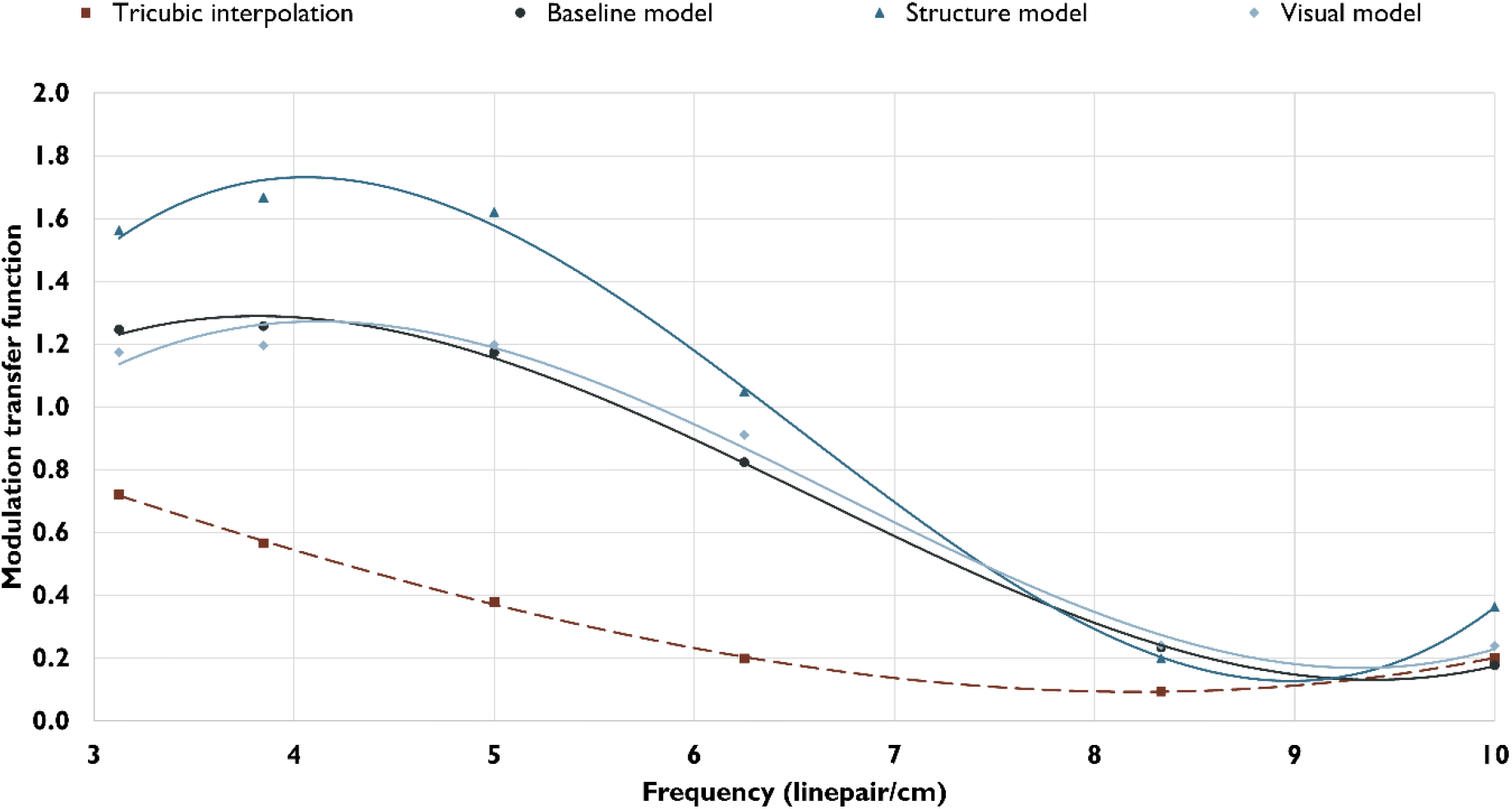
The modulation transfer functions (MTF) are scaled based on the average of plexiglass and water in a region of interest. The super-resolution model’s predictions highlight the structures in the line pair patterns, and the grayscale values exceed the ones in smooth areas of plexiglass. This results in MTF values that exceed one. However, the results also show the effect of highlighting small structures better than the scaling used for Figure 4b.

**Supplementary Figure 2.**
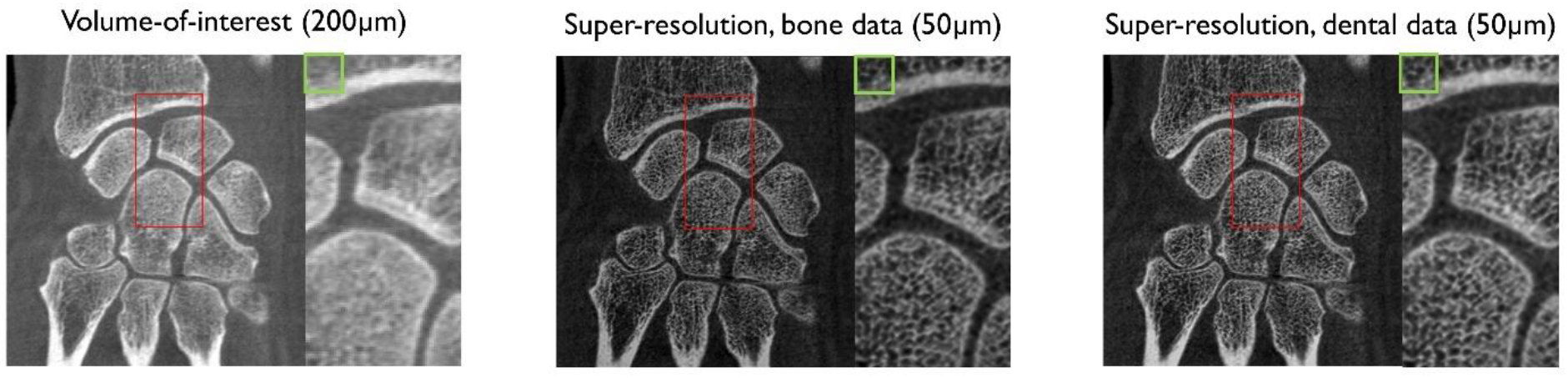
Comparison of using knee tissue blocks and extracted teeth in training data. Structure model predictions are shown above. Only very small differences are seen between the images, suggesting that adding dental images did not improve the prediction accuracy of musculoskeletal cone-beam CT.

**Supplementary Video 1.** Sagittal view of the knee. As the predictions are only created from the transaxial plane, a flickering artefact can be seen on the sagittal view.

**Supplementary Video 2.** Maxillofacial cone-beam CT images of two patients, corresponding structure and baseline model predictions as well as micro-computed tomography (µCT) scans of the extracted teeth. Details are better preserved on the super-resolution prediction. A possible small gap is seen on the lamina dura of patient one, indicated with a red arrow. The tooth next to the tissue is later extracted and the corresponding µCT reconstruction is shown.

## Notes

https://doi.org/10.17632/4xvx4p9tzv.1

https://doi.org/10.5281/zenodo.8041943

### Competing Interest Statement

The authors have declared no competing interest.

## REFERENCES

1. Nieminen MT, Casula V, Nevalainen MT, et al. Osteoarthritis year in review 2018: imaging. Osteoarthritis Cartilage. 2019;27(3):401–411. doi:10.1016/j.joca.2018.12.009

2. Roemer FW, Demehri S, Omoumi P, et al. State of the Art: Imaging of Osteoarthritis— Revisited 2020. Radiology. 2020;296(1):5–21. doi:10.1148/radiol.2020192498

3. Law CP, Chandra R V, Hoang JK, et al. Imaging the oral cavity: key concepts for the radiologist. Br J Radiol. 2011;84(1006):944–957. doi:10.1259/bjr/70520972

4. Roemer FW, Engelke K, Li L, et al. MRI underestimates presence and size of knee osteophytes using CT as a reference standard. Osteoarthritis Cartilage. 2023;31(5):656–668. doi:10.1016/j.joca.2023.01.575

5. Ibad HA, de Cesar Netto C, Shakoor D, et al. Computed Tomography: State-of-the-Art Advancements in Musculoskeletal Imaging. Invest Radiol. 2023;58(1). doi:10.1097/RLI.0000000000000908

6. Segal NA, Li S. WBCT and its evolving role in OA research and clinical practice. Osteoarthritis Imaging. 2022;2(3):100083. doi:10.1016/j.ostima.2022.100083

7. Schulze RKW, Drage NA. Cone-beam computed tomography and its applications in dental and maxillofacial radiology. Clin Radiol. 2020;75(9):647–657. doi:10.1016/j.crad.2020.04.006

8. Vitéz S, Kovács B, Ederer J, et al. Cone beam CT for identifying fractures of the wrist and hand – An alternative to plain radiography? Trauma. 2021;24(3):212–217. doi:10.1177/1460408620984397

9. Veiga C, McClelland J, Moinuddin S, et al. Toward adaptive radiotherapy for head and neck patients: Feasibility study on using CT-to-CBCT deformable registration for “dose of the day” calculations. Med Phys. 2014;41(3):031703. doi:10.1118/1.4864240

10. Zachiu C, de Senneville BD, Tijssen RHN, et al. Non-rigid CT/CBCT to CBCT registration for online external beam radiotherapy guidance. Phys Med Biol. 2018;63(1):015027. doi:10.1088/1361-6560/aa990e

11. Posadzy M, Desimpel J, Vanhoenacker F. Cone beam CT of the musculoskeletal system: clinical applications. Insights Imaging. 2018;9(1):35–45. doi:10.1007/s13244-017-0582-1

12. Brüllmann D, Schulze RKW. Spatial resolution in CBCT machines for dental/maxillofacial applications—what do we know today? Dentomaxillofacial Radiology. 2014;44(1):20140204. doi:10.1259/dmfr.20140204

13. Droege RT, Morin RL. A practical method to measure the MTF of CT scanners. Med Phys. 1982;9(5):758–760. doi:10.1118/1.595124

14. Verdun FR, Racine D, Ott JG, et al. Image quality in CT: From physical measurements to model observers. Physica Medica. 2015;31(8):823–843. doi:10.1016/j.ejmp.2015.08.007

15. Anam C, Fujibuchi T, Budi WS, et al. An algorithm for automated modulation transfer function measurement using an edge of a PMMA phantom: Impact of field of view on spatial resolution of CT images. J Appl Clin Med Phys. 2018;19(6):244–252. doi:10.1002/acm2.12476

16. Friedman SN, Cunningham IA. A moving slanted-edge method to measure the temporal modulation transfer function of fluoroscopic systems. Med Phys. 2008;35(6Part1):2473–2484. doi:10.1118/1.2919724

17. Huda W, Abrahams RB. X-Ray-Based Medical Imaging and Resolution. American Journal of Roentgenology. 2015;204(4):W393–W397. doi:10.2214/AJR.14.13126

18. Ibrahim N, Parsa A, Hassan B, et al. Diagnostic imaging of trabecular bone microstructure for oral implants: a literature review. Dentomaxillofacial Radiology. 2013;42(3):20120075. doi:10.1259/dmfr.20120075

19. Finnilä MAJ, Thevenot J, Aho OM, et al. Association between subchondral bone structure and osteoarthritis histopathological grade. Journal of Orthopaedic Research. 2017;35(4):785–792. doi:10.1002/jor.23312

20. Adams JE. Advances in bone imaging for osteoporosis. Nat Rev Endocrinol. 2013;9(1):28–42. doi:10.1038/nrendo.2012.217

21. Genant HK, Engelke K, Prevrhal S. Advanced CT bone imaging in osteoporosis. Rheumatology. 2008;47(suppl_4):iv9–iv16. doi:10.1093/rheumatology/ken180

22. Chu CR, Williams AA, Coyle CH, et al. Early diagnosis to enable early treatment of pre-osteoarthritis. Arthritis Res Ther. 2012;14(3):212. doi:10.1186/ar3845

23. Karhula SS, Finnilä MAJ, Rytky SJO, et al. Quantifying Subresolution 3D Morphology of Bone with Clinical Computed Tomography. Ann Biomed Eng. 2020;48(2):595–605. doi:10.1007/s10439-019-02374-2

24. He RT, Tu MG, Huang HL, et al. Improving the prediction of the trabecular bone microarchitectural parameters using dental cone-beam computed tomography. BMC Med Imaging. 2019;19(1):10. doi:10.1186/s12880-019-0313-9

25. Kemp P, Stralen J Van, De Graaf P, et al. Cone-Beam CT Compared to Multi-Slice CT for the Diagnostic Analysis of Conductive Hearing Loss: A Feasibility Study. J Int Adv Otol. 2020;16(2):222–226. doi:10.5152/iao.2020.5883

26. Beister M, Kolditz D, Kalender WA. Iterative reconstruction methods in X-ray CT. Physica Medica. 2012;28(2):94–108. doi:https://doi.org/10.1016/j.ejmp.2012.01.003

27. Geyer LL, Schoepf UJ, Meinel FG, et al. State of the Art: Iterative CT Reconstruction Techniques. Radiology. 2015;276(2):339–357. doi:10.1148/radiol.2015132766

28. Thibault JB, Sauer KD, Bouman CA, et al. A three-dimensional statistical approach to improved image quality for multislice helical CT. Med Phys. 2007;34(11):4526–4544. doi:10.1118/1.2789499

29. Greffier J, Frandon J, Si-Mohamed S, et al. Comparison of two deep learning image reconstruction algorithms in chest CT images: A task-based image quality assessment on phantom data. Diagn Interv Imaging. 2022;103(1):21–30. doi:10.1016/j.diii.2021.08.001

30. Szczykutowicz TP, Toia G V, Dhanantwari A, et al. A Review of Deep Learning CT Reconstruction: Concepts, Limitations, and Promise in Clinical Practice. Curr Radiol Rep. 2022;10(9):101–115. doi:10.1007/s40134-022-00399-5

31. Panda J, Meher S. An improved Image Interpolation technique using OLA e-spline. Egyptian Informatics Journal. 2022;23(2):159–172. doi:10.1016/j.eij.2021.10.002

32. Fang L, Monroe F, Novak SW, et al. Deep learning-based point-scanning super-resolution imaging. Nat Methods. 2021;18(4):406–416. doi:10.1038/s41592-021-01080-z

33. Isola P, Zhu JY, Zhou T, et al. Image-to-image translation with conditional adversarial networks. In: Proceedings – 30th IEEE Conference on Computer Vision and Pattern Recognition, CVPR 2017. Vol 2017-January. ; 2017:5967–5976. doi:10.1109/CVPR.2017.632

34. Zhu JY, Park T, Isola P, et al. Unpaired Image-to-Image Translation Using Cycle-Consistent Adversarial Networks. In: Proceedings of the IEEE International Conference on Computer Vision. Vol 2017-October. ; 2017:2242-2251. doi:10.1109/ICCV.2017.244

35. Chaudhari AS, Fang Z, Kogan F, et al. Super-resolution musculoskeletal MRI using deep learning. Magn Reson Med. 2018;80(5):2139–2154. doi:10.1002/mrm.27178

36. Chaudhari AS, Stevens KJ, Wood JP, et al. Utility of deep learning super-resolution in the context of osteoarthritis MRI biomarkers. Journal of Magnetic Resonance Imaging. 2020;51(3):768–779. doi:10.1002/jmri.26872

37. Rudie JD, Gleason T, Barkovich MJ, et al. Clinical Assessment of Deep Learning–based Super-Resolution for 3D Volumetric Brain MRI. Radiol Artif Intell. 2022;4(2):e210059. doi:10.1148/ryai.210059

38. Li H, Prasad RGN, Sekuboyina A, et al. Micro-Ct Synthesis and Inner Ear Super Resolution via Generative Adversarial Networks and Bayesian Inference. In: - 2021 IEEE 18th International Symposium on Biomedical Imaging (ISBI). ; 2021:1500–1504. doi:10.1109/ISBI48211.2021.9434061

39. Yu H, Wang S, Fan Y, et al. Large-factor Micro-CT super-resolution of bone microstructure. Front Phys. 2022;10. doi:10.3389/fphy.2022.997582

40. Tsujioka K, Yamada K, Niwa M. Performance evaluation of micro-vessels imaging by deep learning reconstruction targeting ultra-high-resolution CT (UHR-CT). J Med Imaging Radiat Sci. 2022;53(4, Supplement 1):S28. doi:10.1016/j.jmir.2022.10.093

41. Colbrook MJ, Antun V, Hansen AC. The difficulty of computing stable and accurate neural networks: On the barriers of deep learning and Smale’s 18th problem. Proceedings of the National Academy of Sciences. 2022;119(12):e2107151119. doi:10.1073/pnas.2107151119

42. Johnson J, Alahi A, Fei-Fei L. Perceptual losses for real-time style transfer and super-resolution. In: European Conference on Computer Vision. ; 2016. doi:10.1007/978-3-319-46475-6_43

43. 43. Odena A, Dumoulin V, Olah C. Deconvolution and Checkerboard Artifacts. *Distill*. Published online 2016. http://distill.pub/2016/deconv-checkerboard/

44. Otsu N. A Threshold Selection Method from Gray-Level Histograms. IEEE Trans Syst Man Cybern. 1979;9(1):62–66. doi:10.1109/TSMC.1979.4310076

45. Bouxsein ML, Boyd SK, Christiansen BA, et al. Guidelines for assessment of bone microstructure in rodents using micro–computed tomography. Journal of Bone and Mineral Research. 2010;25(7):1468–1486. doi:10.1002/jbmr.141

46. Menon S, Damian A, Hu S, et al. PULSE: Self-Supervised Photo Upsampling via Latent Space Exploration of Generative Models. In: Proceedings of the IEEE Computer Society Conference on Computer Vision and Pattern Recognition. ; 2020:2434–2442. doi:10.1109/CVPR42600.2020.00251

47. Chaika M, Afat S, Wessling D, et al. Deep learning-based super-resolution gradient echo imaging of the pancreas: Improvement of image quality and reduction of acquisition time. Diagn Interv Imaging. 2023;104(2):53–59. doi:10.1016/j.diii.2022.06.006

48. Van Dyck P, Smekens C, Vanhevel F, et al. Super-Resolution Magnetic Resonance Imaging of the Knee Using 2-Dimensional Turbo Spin Echo Imaging. Invest Radiol. 2020;55(8). doi:10.1097/RLI.0000000000000676

